# Active virus-host system in a Lokiarchaeon culture

**DOI:** 10.64898/2026.02.13.705702

**Authors:** Nevena Maslać, Rafael I. Ponce-Toledo, Thiago Rodrigues-Oliveira, Tomás Alarcón Schumacher, Miroslav Homola, Zhen-Hao Luo, Malcolm F. White, Florian K. M. Schur, Susanne Erdmann, Christa Schleper

**Affiliations:** Department of Functional and Evolutionary Ecology, Archaea Biology and Ecogenomics Unit, University of Vienna, Vienna, Austria; Archaeal Virology research group, Max Planck Institute for Marine Microbiology, Bremen, Germany; Institute of Science and Technology Austria (ISTA), Klosterneuburg, Austria; School of Biology, University of St Andrews, St Andrews, UK; Institute for Microbiology, University of Innsbruck, Innsbruck, Austria

## Abstract

Asgard archaea are considered the closest prokaryotic relatives of eukaryotes and having a crucial role in eukaryogenesis. Only few organisms have been cultivated from this group and their viruses were previously described only through metagenomic reconstructions. Here, we report the first successful cultivation of an Asgard archaeal virus infecting a novel strain of *Ca.* Lokiarchaeum ossiferum B36. The 16 kbp integrated provirus is capable to excise and replicate independently leading to the formation of virus particles. Network analysis of shared protein clusters with other archaeal viruses places it in a new family, *Fylgjaviridae*. The host encodes a distinctive repertoire of antiviral defense systems, including Septu, Wadjet, and type II CBASS system, all different from the defense systems of the related strain *Ca.* L. ossiferum B35. Our cultures provide the first opportunity to study interactions of a virus–host system in Asgards and hold significant potential for developing genetic tools.

## Introduction

Asgard archaea (taxonomically described as Promethearchaeota)^1^ are considered the closest living archaeal relatives of eukaryotes^2^ and represent a huge and widespread group of currently 16 recognized distinct lineages that have been described by genomic and metagenomic approaches^2–4^. More than 390 available metagenome-assembled genomes (MAGs) and 15 closed genomes have been assembled since the original discovery of the first MAGs of Lokiarchaeia, in 2015^5^, according to GTDB^6^ (Nov. 2025). Their analysis revealed that Promethearchaeota harbor several hundred conserved genes encoding what are usually considered essential features of every eukaryotic cell (eukaryotic signature proteins, ESPs)^7,8^. While phylogenetic studies indicate that eukaryotic information-processing systems as well as an actin-based cytoskeleton, ESCRT systems and many other proteins evolved from a common ancestor of Archaea and Eukaryotes^9–12^, it is still unclear whether the same is true for the eukaryotic virome and other mobile genetic elements. The reconstruction of the LECA (Last Eukaryotic Common Ancestor) virome rather indicates a bacterial origin^13^ and there are no plausible candidates for ancestors of eukaryotic viruses among the virus families found in Asgard genomes^14–16^. However, their virome remains largely undersampled and unexplored.

In some of the complete and incomplete Asgard genomes, viruses from different family level groups have been identified, including potentially tailed viruses from the *Caudoviricetes* class^15,16^, putatively icosahedral skuldviruses^14,15^, and wyrdviruses related to spindle-shaped viruses of other archaea^15^. The diversity of viral defense systems detected in Asgard MAGs so far exceeds, that of their virome^17,18^. In total, 2,610 complete viral defense systems have been reported in Asgards, 89 of which are unique to this lineage^17,18^. Notably, among the systems more frequently detected in Asgards than in bacteria or other archaeal groups, viperins and argonauts stand out because of their roles in the evolution of eukaryotic defense systems^17,18^. Conservation of the defence systems in different species can be related to exposure to similar viruses, environmental selection and physicochemical conditions, however, there is no clear correlation between the distribution of defence systems and the latter factors. Overall, their distribution is heterogeneous across the lineages and shaped by extensive loss and gain via horizontal gene transfer^18^, as is evident even when comparing closely related species^19^.

So far, no Asgard archaea viruses have been observed in laboratory cultures, as the host cells themselves are challenging to isolate and sustain cultures. Only two Asgard archaea species have been described so far: *Promethearchaeum syntrophicum*^1,20^ and *Ca.* Lokiarchaeum ossiferum B35^12^, but four more appeared in preliminary announcements^21–23^. All of them share a surprising cellular complexity with cytoskeletal elements^12^, form long protrusions and depend on syntrophic partners, demonstrating that genomic analyses alone can not provide definitive answers to fundamental questions about cell biology and physiology of organisms. Similarly, intricate contacts and dynamics between viruses and host cells can not be inferred from genomes only.

Here, we present the first stable host–virus system of an Asgard archaeon in culture, consisting of a novel strain of *Ca. Lokiarchaeum ossiferum* (B36) and its active provirus, *Fylgjavirus 1*, a member of the viral class *Tectiliviricetes*, which includes several families of viruses infecting members of all three domains. Combining cultivation approaches with bioinformatics we characterized this novel virus-host system and analyzed the defensome of both Ca. *L. ossiferum* strains, revealing surprising differences and underscoring the role of horizontal gene transfer in shaping it.

## Results

### Enrichment of *Ca.* Lokiarchaeum ossiferum strain B36

We utilized sediment of an estuarine canal in Slovenia as inoculum for cultures subjected to different growth conditions, similar to the previously described enrichment of *Ca.* Lokiarchaeum ossiferum strain B35^12^ (Loki-B35). However, in contrast to the previous enrichments of Loki-B35, where the sea water from the sampling site was used as a base medium, here we inoculated the sediment slurries directly to the MKD-1 medium^20^. During the subsequent transfers, the cultivation conditions were matched to those of Loki-B35^12^ (see Material and Methods). However, in comparison to Loki-B35, which reaches cell densities of up to 5 × 10^7^ cells/mL in enrichments ranging from 24 to 79% lokiarchaeal relative abundance, these cultures displayed notably lower values of up to 1.5 × 10⁶ cells/mL and 10 to 37% relative enrichments (Figure 1a). A single circularized lokiarchaeal genome was assembled from this enrichment and revealed an average nucleotide identity of 97.2% with the genome of *Ca.* L. ossiferum B35. Furthermore, a comparison of the sequence identity among the 3 copies of 16S rRNA genes present in each genome showed a range between 99.7% to 99.9%. Based on established species and strain boundaries for prokaryotic genomes^24^, these findings indicate that this organism constitutes a new strain within the same species, henceforth named *Ca*. L. ossiferum strain B36 (Loki-B36) (Figure 1b). Amplicon sequencing of prokaryotic 16S rRNA genes revealed four main groups in the enrichment: a single Lokiarchaeon (30.2%), a sulfate-reducing bacterium of the *Halodesulfovibrio* genus (32.4%), a bacterium of the *Izemoplasmataceae* group (32.4%), a methanogen from the *Methanosarcina* group (2.5%), and few other microbial groups of low abundance (2.5%) (Figure 1c). It is noteworthy that both previous lokiarchaea enrichments of *P. syntrophicum* and Loki*-*B35 also included a sulphate reducer and a methanogen^12,20^. Near complete MAGs (>98%) of the *Halodesulfovibrio* partners were assembled from B35 (Halo B35) and B36 (HaloB36) enrichments as well as the genome of the *Izemoplasmataceae* bacterium present in B36 cultures, their genomic characteristics are shown in Supplementary Table 1.

**Fig. 1:**
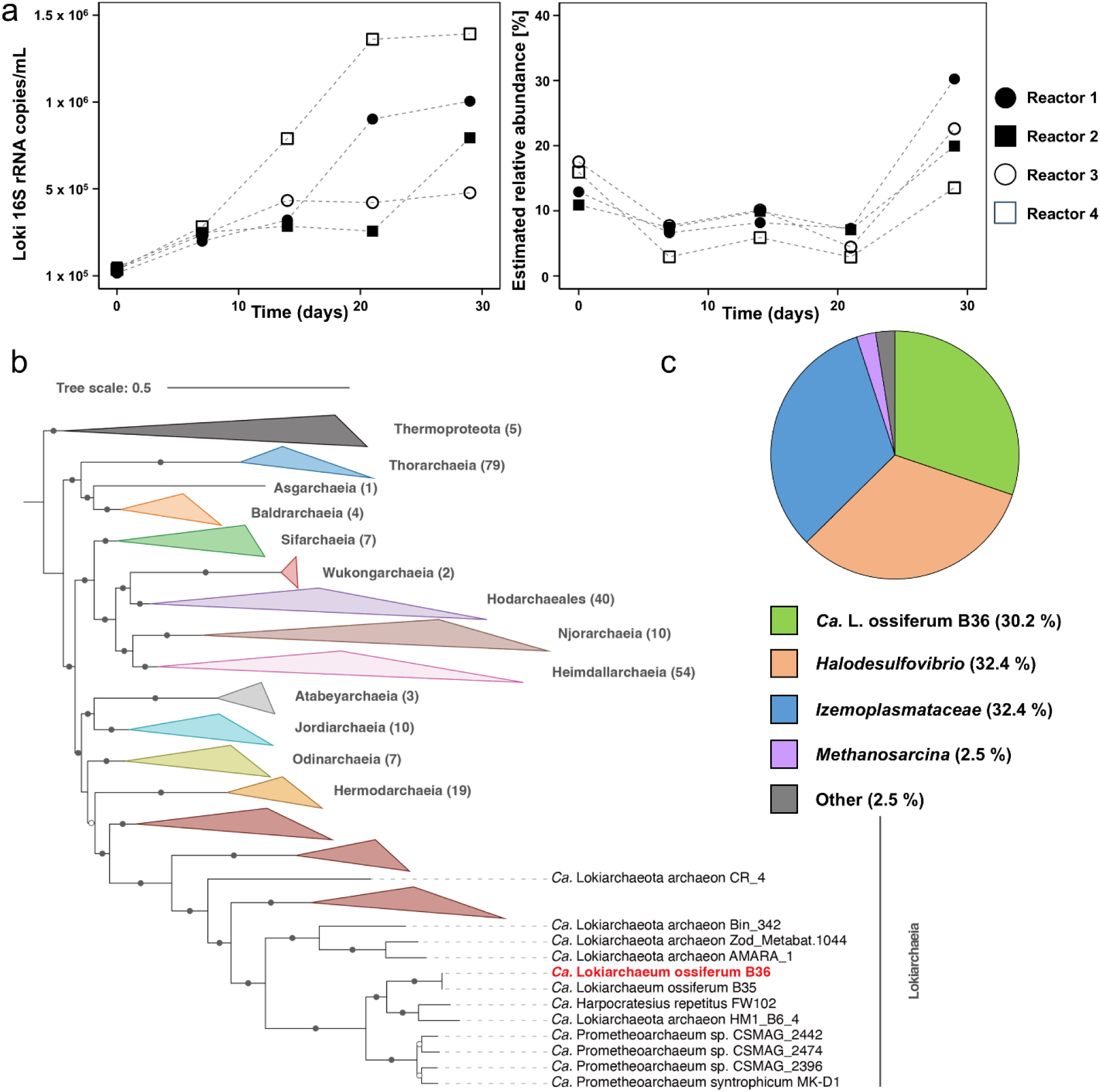
Growth, enrichment and the phylogenetic placement of *Ca.* Lokiarchaeum ossiferum B36. **a,** Loki-B36 reaches a maximal cell density of 1.39 × 10^6^ /mL and a relative abundance of 37.17% in a 2.2 L reactor system (MLM, 80:20 N_2_:CO_2_, n=4). Growth was quantified by qPCR (see Material and Methods). **b,** Maximum-likelihood (ML) phylogeny of *Ca.* L. ossiferum B36 based on archaeal marker genes identified by GTDB-Tk (see Methods). *Ca.* L. ossiferum B36 represents a new strain within the *Ca.* L. ossiferum species. **c,** Loki-B36 enrichment composition determined by 16S rRNA gene amplicon sequencing.

*Ca.* L. ossiferum B36 displayed the largest genome of all other closed archaeal genomes described to date, with over 6.2 Mb and 5 332 predicted proteins (Figure 2a). Synteny analysis between the two strains revealed a total of 12 medium-to-long-range genomic rearrangements (Supplementary Figure 1). Comparative analysis of protein families between both Loki strains revealed that they share a total of 3 948 proteins (Figure 2b), illustrating a high degree of similarity in protein content. Among the clusters that could be functionally annotated, differences between Loki-B35 and Loki-B36 were mostly noted in proteins associated with carbohydrate transport, defense mechanisms, and the mobilome (Figure 2c, Supplementary Tables 3 and 4). Notably, Loki-B36 contained more virus-associated protein families (related to defense systems and mobile elements), as well as proteins involved in translation, ribosomal structure and biogenesis, while Loki-B35, is enriched in proteins involved in amino acid and carbohydrate transport and metabolism.

**Fig. 2:**
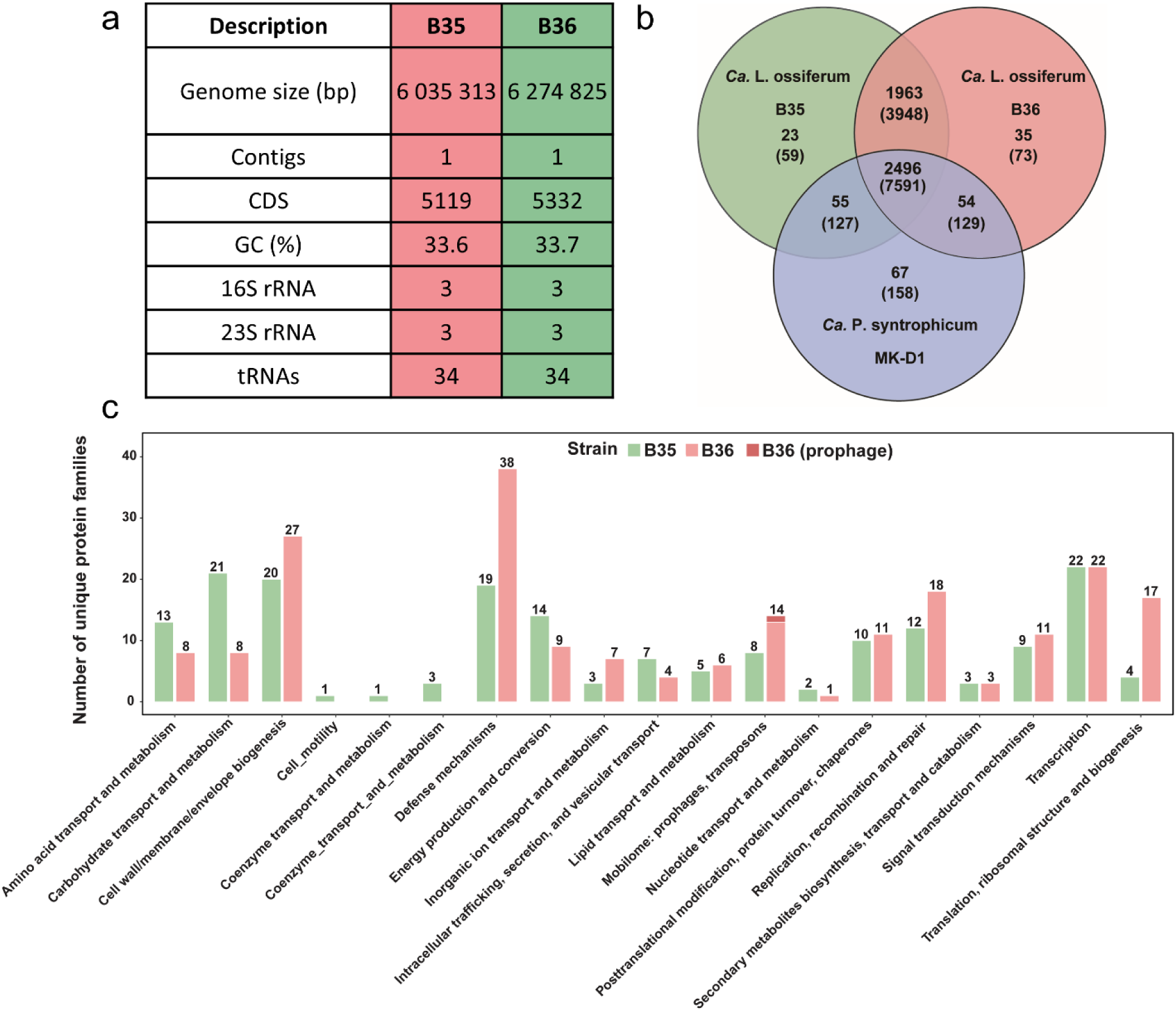
Genome analysis of *Ca.* Lokiarchaeum ossiferum B36. **a,** Characteristics of the genome of *Ca.* L. ossiferum strain B36 in comparison to strain B35. **b,** Number of shared protein families between strains B36, B35 and *Promethearchaeum syntrophicum*. For each genome, the number above the parentheses represents the number of protein clusters, while the number inside the parentheses corresponds to the total number of proteins. **c,** Number of unique proteins in different functional categories in the genomes of Loki-B35 and B36. Annotation was performed according to the asCOG database2 (see Material and Methods).

Scanning electron microscopy (SEM) analysis showed that, akin to strain B35, strain B36 exhibits a discernible round cell body with extensive protrusions of variable sizes, occasional branching, and the presence of bulbous structures along their length and at the ends (Figure 3a-b). In addition to the single cells in liquid cultures, close contact among individual lokiarchaea cells and also with syntrophic partners was frequently observed by SEM imaging (Figure 3c-d). Even very large clusters of cells of up to 20 µm in diameter were regularly seen (Figure 3c-d). The similarities of both strains extend to their ultrastructural organization, as revealed by cryo-electron tomography (cryo-ET) analysis (Figure 3e). As reported earlier for the B35 strain, for Loki-B36 we also observed a highly pleomorphic cell shape with a cell body and protrusions containing numerous ribosomes, a single plasma membrane lacking an S-layer, but instead showing unordered surface densities (Figure 3f), as well as thin protein densities connecting across membranes (e.g. from the cell body to protrusions) (Figure 3e). Cytoskeletal filaments resembling the Lokiactin filaments were also observed in cryo-electron tomograms (Figure 3e).

**Fig. 3:**
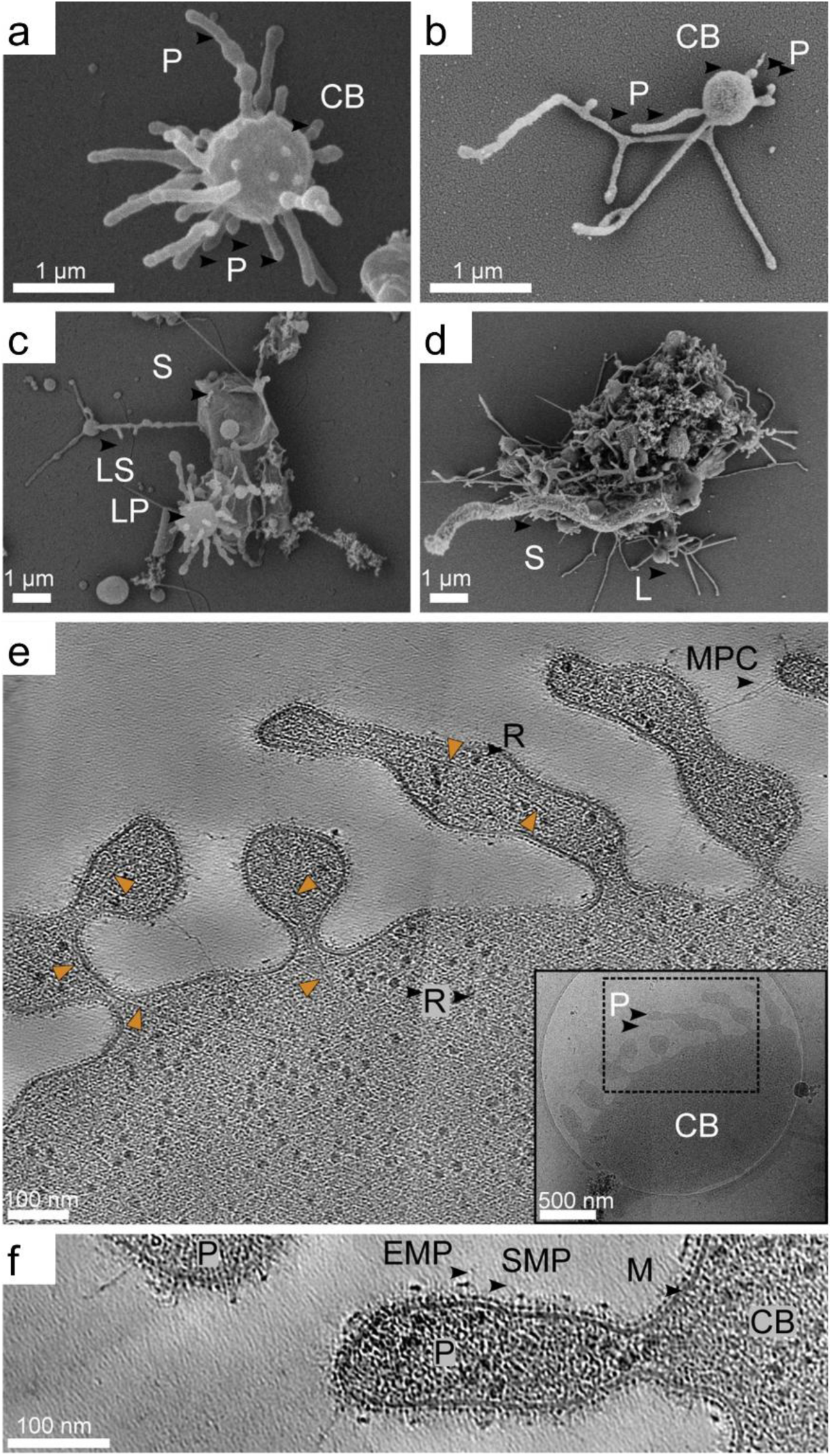
Morphology and ultrastructure of *Ca.* L. ossiferum B36 strain cells. **a-d,** SEM imaging of fixed Loki cells. **a,b,** Single small coccoid cells with extensive protrusions in planktonic form (**a**) and settled form (**b**). CB, cell body; P, protrusion. **c,d.** Loki cells often create a cluster with co-cultured organisms. L, Loki cell; LS, Loki settled cell; LP, Loki planktonic cell; S, symbiont cell. Scale bar, 1 μm. **e,f,** Slices through denoised cryo-tomograms, 14 nm thick, of near-native Loki cells. **e**, Edge of the Loki-B36 cell body with extensive protrusions containing Lokiactin filaments (orange arrowheads), ribosomes, R, and displaying membrane protein connection, MPC. Scale bar 100 nm. Inset shows overview projection micrograph of the target cell displaying its typical morphology with central body, CB, and protrusions, P. Scale bar 500 nm. **f,** Magnified view on the Loki protrusion displaying several layers of membrane proteins. M, membrane; SMP, surface membrane proteins layer; EMP, envelope membrane proteins.

### Loki-B36 encodes an active virus

In a genomic region that was conserved in both strains, a putative provirus of 16.1 kb was found inserted in strain B36 (Figure 4a). In only one out of six sequenced metagenomes of enrichment cultures, the provirus was discovered as a genomic region with significantly higher read coverage (approx. 6-8 fold) relative to the rest of the genome, indicative of increased extrachromosomal replication in this specific region and suggestive of spontaneous virus production (Figure 4a, lower panel).

**Fig. 4:**
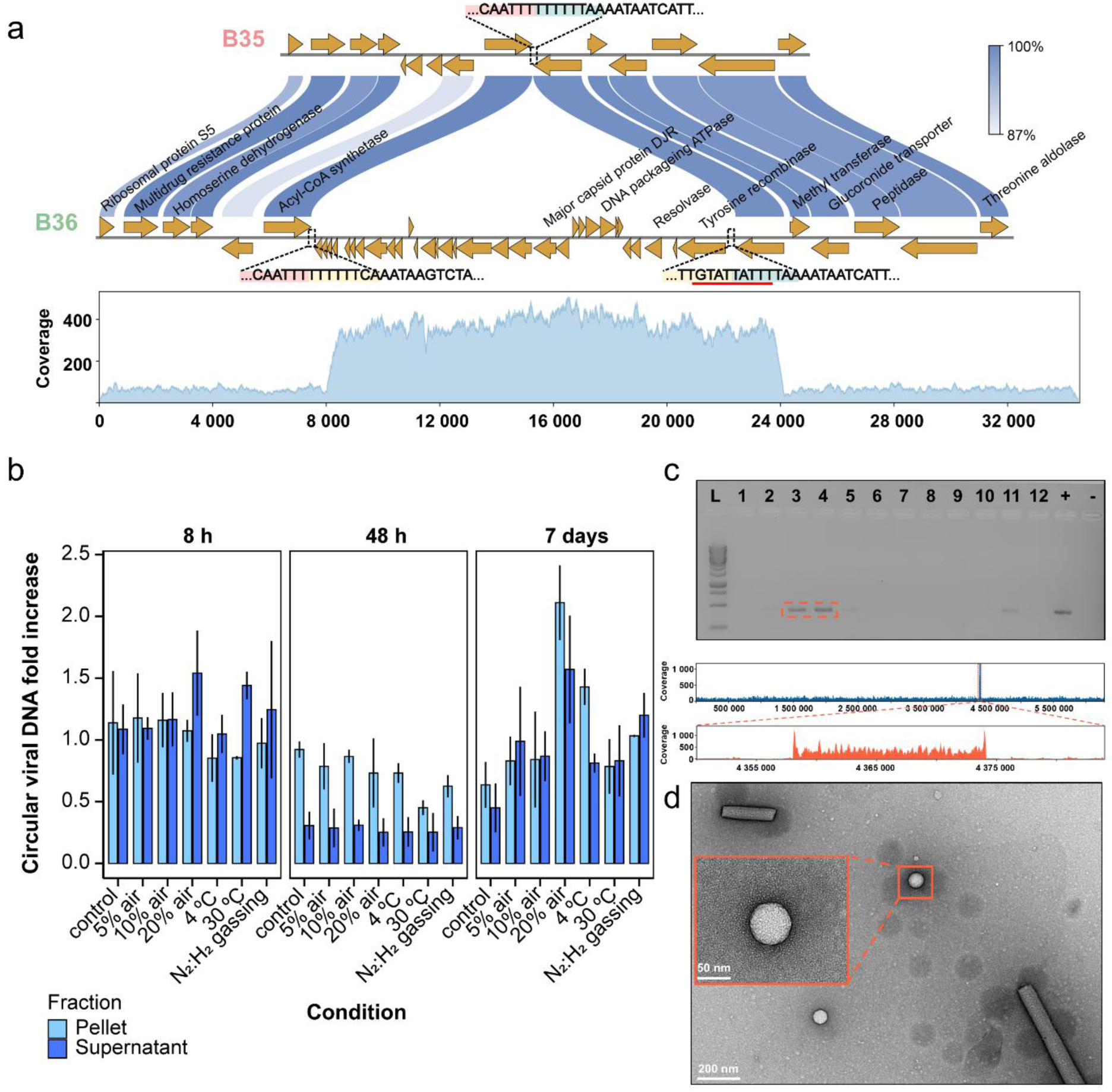
Characterization, induction and purification of Fylgjavirus 1. **a,** Gene synteny comparison between *Ca.* L. ossiferum B35 and B36 and putative viral region with high coverage detected in strain B36. Bar shows the percentage of similarity between proteins. Highlighted sequences are color-coded as following: pink/yellow and yellow/blue: Fyl1 integration sites, red underline: inverted repeat. **b,** Influence of different stress conditions on viral particle production in Loki-B36 (n=3). The average increase was calculated by dividing the values obtained from each replicate with an average from the control calculated at time point zero. The number of viral particles were quantified by qPCR in both cell pellets and culture supernatant (see Methods). **c,** Screening of the CsCl gradient fractions from the virus purification for the presence of the virus by PCR (L=ladder, 1-12 corresponding CsCl fractions, + positive control, - negative control) shown in an agarose gel. DNA from fraction 3 and 4 were subjected to Illumina sequencing. Coverage of the B36 genome with a peak corresponding to the provirus genome, and a zoom onto the provirus region, is shown. **d,** Negative staining electron micrograph of FylV1 candidate particle: left: isosahedral virus-like particle with a diameter of ∼ 55 nm (scale bar 50 nm) and its environment (scale bar 200 nm).

To test whether viral particles could be detected in cultures, we first screened both cell pellets and culture supernatants of B36 cell cultures at various time points by qPCR, using primers targeting the circularized viral genome, (Supplementary Figure 2a). We detected an initial decline of circularized virus genome copy number per cell upon inoculation of a fresh culture. However, as the cells grew, the virus replicated, leading to a steady increase in the number of circular virus genomes per cell. The circular virus-to-host genome ratio remained low (max. 0.15), and never reached the numbers initially detected in one of the cultures by genome sequencing (Supplementary Figure 2a). During the later stages of cell growth, a higher number of active viral particles were released into the supernatant (Supplementary Figure 2b), though still at very low numbers (approximately 10^5^ particles/mL). Therefore, we assume that the virus is only active (producing virus particles) in a small fraction of the population, as no significant decline of Loki-B36 cell numbers was observed.

### Infection and Induction experiments with the Loki-B36 virus

To test, if the virus of strain B36 could infect the closely related strain B35, which also contains an almost identical homing site in its genome (Fig. 4a), we used both directly filtered B36 cultures in the exponential growth phase as well as their supernatants and mixed it with growing B35 cultures in different amounts (ranging from 0.1 to 10% of the culture volume, Supplementary Figure 3). As a control, B35 culture mixed with the B35 supernatant was used. However, no indication of a successful infection was obtained (Supplementary Figure 3).

To search for conditions under which the virus might be induced in its host B36, the cultures were subjected to various stress conditions, including introduction of different amounts of atmospheric air in the gas phase (5%, 10%, 20%), cold (incubation at 4 °C) and heat shock (incubation at 30 °C), and addition of N₂:H₂ mixture (75:25%) to the gas phase (Figure 4b). Circular virus DNA was quantified in both the cell pellet and the culture supernatant over a period of up to 7 days (Figure 4b). The highest fold increase in viral genomes, i.e. twofold, was observed after 7 days of incubation in cultures where 20% of atmospheric air was added to the gas phase (Figure 4b). In all other conditions, viral numbers did not increase by more than 1.5-fold. Subsequently, we tested prolonged incubation times with lower air percentages (5% and 10%) and monitored the circularized viral DNA over a period of three weeks (Supplementary Figure 4). In this case, the addition of 5% atmospheric air to the gas phase again produced the stronger effect, but the level of induction remained unchanged at twofold (Supplementary Figure 4). Induction through other conventional triggers, such as the addition of mitomycin C, UV stress and nutrient depletion, did not yield increased virus production (Supplementary Table 5).

To identify virus particles, we purified concentrated culture supernatants by CsCl density centrifugation. Although particle bands were not visible in gradients, the virus genome was detected in three fractions of the gradient by PCR (Figure 4c). Illumina sequencing of the DNA isolated from these bands, allowed the full assembly of the circular virus genome (mean coverage approx. 400x), with only minor host contaminations (average coverage, excluding the provirus, approx. 20) (Figure 4c). Purification by CsCl gradient eliminated some, but not all non-viral structures observed while imaging concentrated culture supernatants with a transmission electron microscope (TEM). We observed several virus-like particle (VLP) candidates (Figure 4d). However, we could not identify them based on their shape, as they were not uniform enough to sort into a strong individual class upon 2D-classification.

### A virus of a new family: ‘*Fylgjaviridae’*

The provirus present in B36 has a 16.1 kb genome (16 070 bp) with an inverted repeat at the end and encodes for 32 putative proteins. Contrary to other described proviruses in Asgard genomes, and many other archaeal and bacterial systems often found in tRNA genes^25^, the provirus in B36 was found integrated between a gene coding for an acetyl-coenzyme A synthetase (AMP-forming), and a gene coding for a hypothetical protein with armadilllo (ARM) repeats (spanning between positions 4 358 897 and 4 374 967 on the Loki-B36 genome, Figure 5a). The B36 virus encodes a double jelly-roll (DJR) major capsid protein (MCP), which is characteristic for viruses belonging to the *Varidnaviria* realm^26^ and commonly associated with icosahedral morphology^26,27^. Members of this group infect organisms across the whole tree of life, and related varidnaviruses from the *Skuldviridae* family have been previously predicted by metagenomics to infect members of the Lokiarchaeia^15^. However, genome comparison revealed low levels of similarity with other skuldviruses (Figure 5a), with only the MCP and two hypothetical proteins displaying significant similarity (26%, 27% and 29% respectively, in order), while also lacking the characteristic colinearity of skuldviruses. Furthermore, the virus also lacks the proposed replication protein of skuldviruses^15^.

**Fig. 5:**
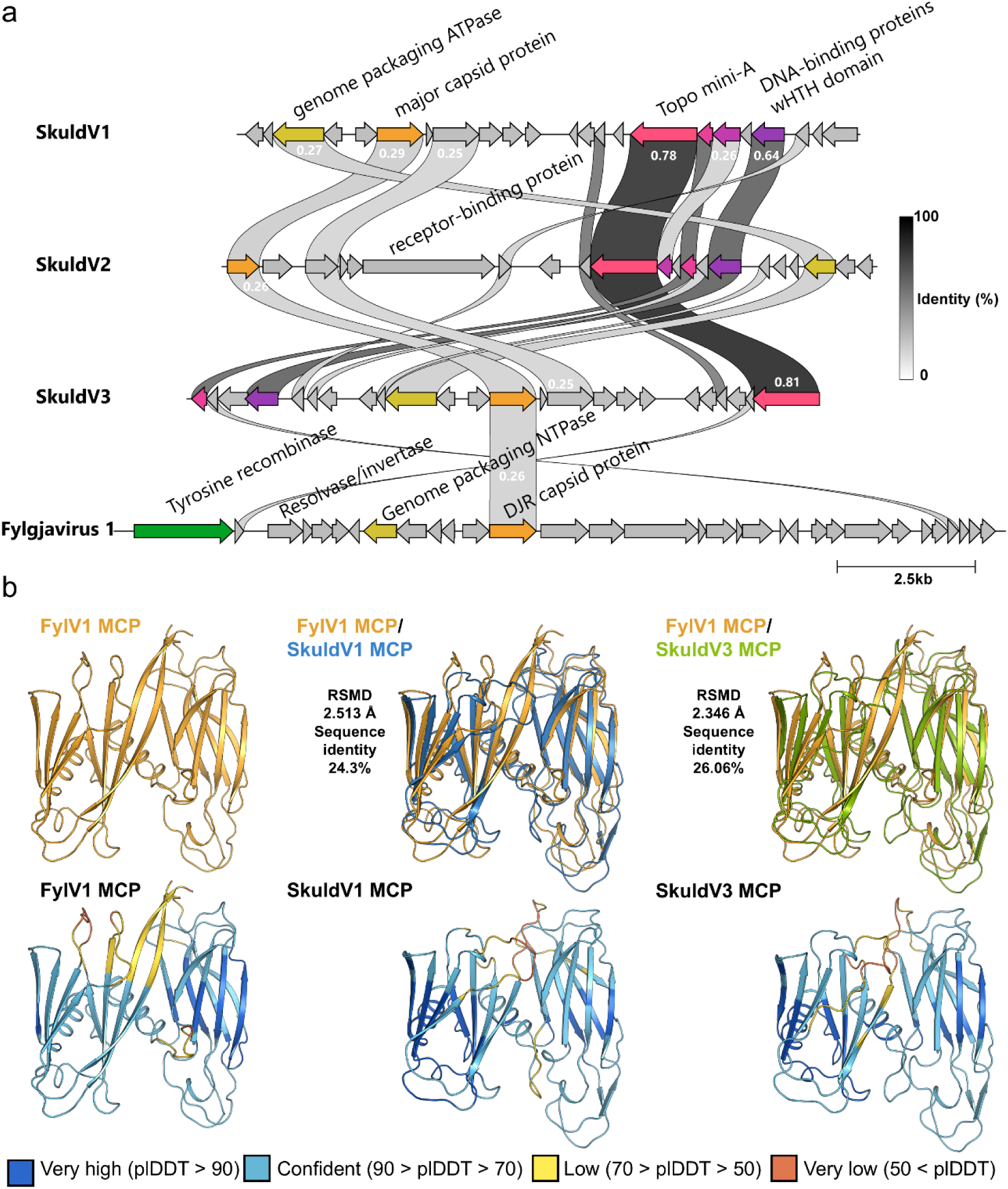
Genome analysis of Fylgjavirus 1. **a,** Genomic alignment of related Skuldviruses and Fylgjavirus 1. Similarity values between proteins (blastp) are indicated in grayscale. Homologues of conserved genes are color coded as follows: major capsid protein (orange), genome packaging ATPase (yellow), Topo mini-A (magenta), ATPase (magenta), DNA-binding proteins (lilac), and Tyrosine recombinase-like protein (green). **b,** Structural prediction and comparison against selected MCPs, of the major capsid protein (MCP) of the Fylgjavirus 1 generated with AlphaFold2. AlphaFold 2 models are shown using ribbon representation and color-coded as follows: top: FylV1 MCP; sky blue SkuldV1 MCP, cyan SkuldV3 MCP; bottom: all three models are color-coded according to their pLDDT score representing per atom confidence estimate on a 0-100 scale: dark blue = very confident (pLDDT > 90), light blue = confident (90 > pLDDT > 70), yellow = low (70 > pLDDT > 50), orange = very low (50 < pLDDT).

By Hidden Markov Model (HMM) comparison, the MCP of the B36 virus appears closely related to the *Turriviridae* family (Sulfolobus turreted icosahedral virus 1-STIV1-; probability = 98.1%), the *Corticoviridae* (Pseudoalteromonas phage PM2; probability = 97.73%) and *Tectiviridae* (Enterobacteria phage PRD1 probability = 94.23%). Additionally, it is distantly related to the eukaryote-infecting Paramecium bursaria Chlorella virus 1 MCP (probability = 81.72%). Structural prediction of the MCP performed with AlphaFold^28,29^, and further structural comparison against other reported archaeal virus MCP structures using Dali^30^, showed that the MCP of the B36 virus is most closely related to SkuldV1 and SkuldV3 viruses (Figure 5b). These two viruses were discovered through metagenomics and associated with asgardarchaeal hosts^15^, and form together with the B36 virus a clade distinct from the one containing STIV1^31^ (Supplementary Table 6).

Functional annotation using protein sequences and predicted structures suggests that the virus genome is likely organized in a modular form. The structural module contains the MCP and other accessory proteins with low levels of similarity to phage structural proteins (ORFs 13 to 20, Supplementary Table 6). Protein analysis of purified STIV virions revealed several accessory minor structural proteins^31^, further suggesting that the predicted ORFs could play important roles in the virion structure. Furthermore, similar to most varidnaviruses, the B36 virus encodes a genome packaging NTPase that also displays high levels of structural homology with the one from STIV2 (PDBid 4kfr, sequence identity = 19.5; TM-Score= 0.74615 and RMSD = 3.29).

Interestingly, while corticoviruses replicate through the rolling-circle mechanism, using a HUH-like endonuclease^32^, and skuldviruses are hypothesized to replicate using a standalone topoisomerase-like protein called Topo Mini-A^15,33^, no homolog for these or any other known replication protein were identified in the B36 virus genome.

Therefore, we propose that the B36 virus represents a separate virus family within the *Tectiliviricetes* order, which we propose to name *Fylgjaviridae* (in Nordic mythology, a fylgja is a supernatural being or spirit which accompanies a person in connection to their fate or fortune. The word fylgja means, “to accompany”), with the B36 virus, being currently the only member, named Fylgjavirus 1 (FylV1).

### The two *Ca.* L. ossiferum strains have distinct viral defense systems

With 6 defense systems in Loki-B35 and 8 defense systems in Loki-B36 (Figure 6, Supplementary Figure 5, Supplementary Table 7), both strains have a slightly higher defense system–per–genome ratio than other Promethearchaeota (average 4.9) and, more specifically, the Lokiarchaeia (average 5.8 defense systems per genome)^17^. In contrast to *P. syntrophicum* (Supplementary Figure 6), the genomes of *Ca.* L. ossiferum strains do not contain CRISPR arrays. Instead, they harbor a number of antiviral defense systems that are remarkably distinct among the two closely related strains, although both were enriched from the same habitat (Figure 6, Supplementary Figure 5, Supplementary Table 7). Strain Loki-B35 encodes a prokaryotic Viperin (pVip), a Restriction-Modification (R-M) system with a putative DrmD component and two Pycsar systems that function via cUMP or cCMP signaling (detected by PADLOC^34,35^). The Pycsar systems cluster with an operon encoding Gabija – a two-protein system that recognizes and cleaves invading DNA^19,36^. On the other hand, strain Loki-B36 lacks Viperin, Gabija and Pycsar systems and instead encodes the Septu antiviral nuclease complex^19^, the Wadjet anti-plasmid SMC family ATPase^19,37^ and a type II CBASS system that activates programmed cell death upon detection of viral infection^38^. Among these, CBASS, Mokosh and viperins are unique to Asgards^17^. The more distant lokiarchaeon *P. syntrophicum* displays yet another combination of defense systems: R-M, Gabija, pVip, SoFic, Mokosh type II and CBASS type II. Interestingly, we detected almost all antiviral defense proteins from Loki-B35 in the proteome except for pVip, while in the case of Loki-B36, less than half of the proteins from defense systems were observed (Supplementary Table 7).

**Fig. 6:**
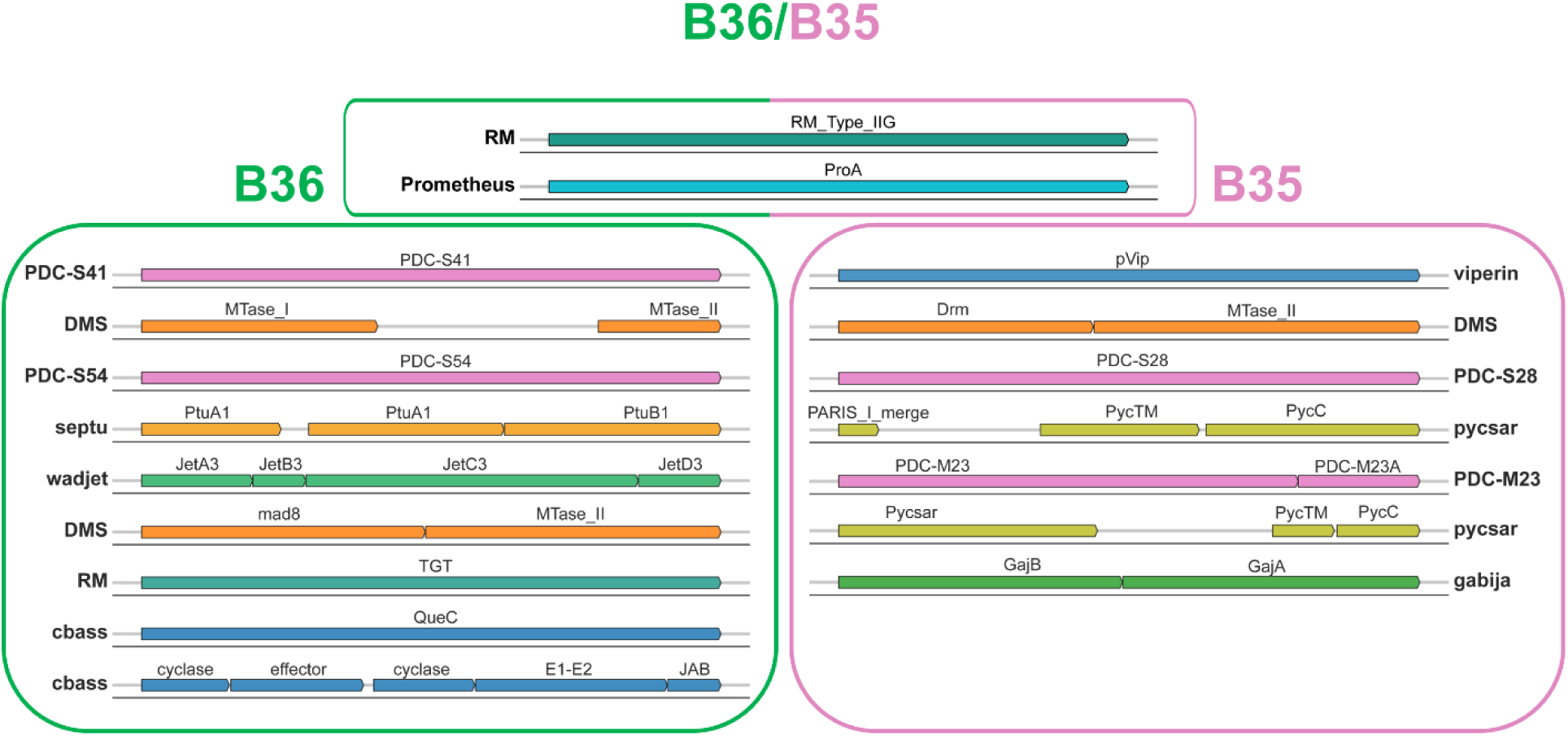
Comparison of antiviral defense systems of both strains of *Ca.* L. ossiferum. The two strains B35 and B36 share only two defence systems: RM Type IIG and Prometheus. The genes are drawn to scale, however the information about the position in the genome is left out for simplicity and displayed in Supplementary Figure 5.

Furthermore, to investigate whether any of these defense system proteins have co-evolved with their host lineages, we compared the phylogenetic topologies of individual gene trees with the Asgard species tree (Supplementary Figure 7). A topological comparison analysis revealed that gene trees of all three defense systems shared among both Loki strains (Prometheus, RM, DMS) displayed significantly greater congruence with the species tree than expected by chance (Supplementary Figure 7), indicating vertical inheritance as expected.

The phylogenetic analyses of DrmD, a SNF-2 like helicase encoded in the genome of B35, and the CBASS system present in B36 and *P. syntrophicum* showed that these genes were likely transferred from a bacterial donor (Supplementary Figure 8.)

## Discussion

*Ca.* L. ossiferum strain B36 and FlyV1 represent the first virus-host system of an Asgard archaeon available in culture, providing insights into virus-host interactions in this lineage.

FlyV1 is a temperate provirus producing virus particles only in a small fraction of host cells, suggesting a preference for a propagation in a rather passive form (vertical) with its host (piggyback-the-winner as opposed to kill-the-winner relationship). Low virus-to-host ratios have been described previously, for example in extreme environments and the human gut^39,40^. Most described members of the *Varidnaviria* realm are lytic viruses, and with the recently described skuldviruses of Asgard archaea thought to be lytic as well^15^, we propose that FlyV1 exhibits a lytic life cycle. Considering the relatively long doubling time of the host, we suggest that a low virus-to-host ratio is likely beneficial for both host and virus, allowing a stable number of free virus particles without dramatically reducing host cell numbers while possibly protecting the host from other viruses.

Viral infections are a major determinant of microbial community compositions, especially in the aquatic ecosystems^41^. Although data specifically on archaeal communities are lacking, up to 40% of the marine prokaryotes are estimated to be lysed by viruses daily^41,42^, making viruses the primary cause for mortality. This, in turn, creates immense evolutionary pressure for a flexible defensome, reflected in the high rates of horizontal gene transfer of viral defense system genes, and resulting in their patchy distribution^18^, causing even closely related strains to possess very different viral defense repertoires, as seen in Loki-B35 and B36, as well as in *P. syntrophicum*.

Like the Asgard viruses recently retrieved from Asgard MAGs^14–16^, FlyV1 is not closely related to eukaryotic viruses and instead takes up a unique position within the virosphere, in line with the observations that the majority of archaeal viruses are exclusive to their hosts and belong to distinct evolutionary lineages^43^. This, however, does not exclude the exciting prospects of discovering novel Asgard archaeal virus groups with roles in the evolution of eukaryotic viruses.

Our work provides first insights into an Asgard virus - host system and will allow deeper insights into their interaction in the future. Having a virus-free closely related strain (B35) in the laboratory also opens the door for the development of a genetic system for Asgard archaea based on transfection experiments with viral DNA.

## Methods

### Sample collection and *Ca*. L. ossiferum B36 strain enrichment

Sediment core samples were retrieved from a shallow estuarine canal near the coast of Piran, Slovenia on April 21^st^, 2019, and processed in an anaerobic tent as previously described^12^. Like in the *Ca.* L. ossiferum strain B35 enrichment process, the layer with the highest amount of lokiarchaeal 16S rRNA gene copies in relation to total DNA content was selected for cultivation in 120 mL serum bottles sealed with butyl rubber stoppers, using 2 g of sediment as inoculum. DNA extraction was performed using the NucleoSpin Soil DNA extraction kit (Macherey-Nagel). qPCR assays using primers LkF (5′-ATC GAT AGG GGC CGT GAG AG) and LkR (5′-CCC GAC CAC TTG AAG AGC TG) targeting lokiarchaeal 16S rRNA genes were performed as previously described^12^ and normalized for the number of 16S rRNA copies (three) and extraction volume. After 190 days, growth was detected in flasks containing 50 mL of MK-D1 medium^20^ and incubated at 20 °C (0.3 bar of N_2_:CO_2_ (80%:20%:)) supplemented with ampicillin, kanamycin and streptomycin (50 µg/mL each). After three transfers in these conditions, the medium was changed, and the rest of the cultivation was performed in batch, in MLM medium as previously described^12^.

### Cultivation in 2.2 L bioreactor setup

*Ca*. L. ossiferum B36 was also cultivated in the Eppendorf DASGIP® Bioblock reactor system, equipped with 4× 2.2 L Bioblock stirrer reactors (Eppendorf AG, Hamburg, Germany) with a working volume of 2.0 L (reactor culture). The cells were cultivated without stirring, with the gas flow of 10 sL/H (N_2_:CO_2_, 80:20 %) at 20°C, in MLM medium. As inoculum, cells grown in batch culture up to exponential phase were used in a ratio of 1:10. The absolute numbers of *Ca*. L. ossiferum B36 in the samples were determined via qPCR as described above. The relative abundance of *Ca*. L. ossiferum B36 was estimated based on the number of total genome copies per 1 ng of extracted DNA calculated using the average base pair weight for double stranded DNA.

### 16S rRNA gene amplicon sequencing

The general prokaryotic 16S rRNA gene targeting primer pair 515F (5’-GTG CCA GCM GCC GCG GTA A) and 806R (5’-GGA CTA CHV GGG TWT CTA AT)^44^ was used in PCR assays using lokiarchaeal enrichment culture DNA as template. The resulting amplicons were barcoded and sequenced at the Vienna BioCenter Core Facilities (VBCF) using the Illumina Miseq (300 PE) platform. Cutadapt^45^ was used to remove primer sequences and the QIIME2 pipeline^46^ was used for sequence analysis. The DADA2 algorithm was used to denoise the data and remove low-quality reads and chimeras. Sequences with 100% sequence identity were clustered into amplicon sequence variants (ASV). Taxonomy of ASVs was assigned using the SILVA database (release 138) with the “q2-feature-classifier” plugin^47^.

### Metagenomic sequencing and genome assembly of B36 genome

Illumina sequencing of three samples from our B36 enrichment was performed using the NovaSeq 6000 (paired-end, 150 bp) platform at Novogene. Sequencing data were processed using Trimmomatic (v.0.36)^48^ to remove Illumina adapters and low-quality reads (SLIDINGWINDOW:5:20 LEADING:5 TRAILING:5 MINLEN:50 HEADCROP:6). For long-read metagenomic sequencing, library preparation was carried out with the SQK-LSK109 ligation kit followed by the sequencing in the high-throughput nanopore PromethION platform using FLO-PRO002 flowcells. The fast5 files were basecalled using the high-accuracy ONT basecaller Bonito (v.0.3.6; https://github.com/nanoporetech/bonito) with the dna_r9.4.1 pretrained model (bonito basecaller --fastq dna_r9.4.1). Adapter removal and long-read demultiplexing was carried out with Porechop (v.0.2.4)^49^. NanoFilt (v.2.8.0; NanoFilt −l 5000)^50^ was used to remove reads shorter than 5 Kb. Metagenomic assembly was performed using a similar approach to the *Ca.* L. ossiferum B35 genome^12^. Briefly, long reads were assembled using metaFlye (v.2.8.3-b1695)^51^ producing a circular lokiarchaeal genome that was polished with short reads using four rounds of Pilon^52^ as part of the previously proposed validation pipeline of metagenomic assemblies^53^.

### Comparative genomics

Genome annotation and protein prediction of *Ca.* L. ossiferum B36 was performed using Prokka (v.1.14.6)^54^. To conduct additional functional annotation of genes, the orthology numbers of Kyoto Encyclopedia of Genomes (KEGG) were retrieved using the BlastKOALA online server^55^ and protein domain families were identified using HMM searches against the Pfam database (v.34.0)^56^. The asCOG (Asgard clusters of orthologous genes) annotation of *Ca.* L. ossiferum B36 proteins was performed as previously described.^12^. Protein clustering of orthologous genes between *Ca.* L. ossiferum B35, *Ca.* L. ossfierum B36 and *P. syntrophicum* was carried out using OrthoVenn2^57^. Singletons were considered as species-specific clusters. Average nucleotide identity (ANI) between *Ca.* L. ossiferum B35 and B36 was calculated with the FastANI tool v1.33^24^. The identification and annotation of defense systems was carried out using the PADLOC v2.0.0 tool^34^ and the results can be found in Supplementary Table 7.

### *Ca.* L. ossiferum B36 proteome

*Ca.* L. ossiferum B36 enrichments cultures at exponential growth phase were centrifuged at 20 000 x *g* for 30 min at 4 °C. The resulting pellet was frozen in liquid nitrogen and stored at −80 °C until further processing. Proteome preparation and analysis were performed using the services of the Mass Spectrometry Facility at Max Perutz Labs in Vienna, Austria, using the VBCF instrument pool, as described previously^58^.

### Circularized viral genome detection

*Ca.* L. ossiferum B36 cultures were centrifuged at 20 000 x *g* for 30 minutes, 4 °C, and both the pellet and supernatant fractions were used for DNA extraction. To remove cellular contaminations, the supernatant was filtered through 0.45 µm cellulose membranes (Whatman) and concentrated on Amicon Ultra-15 Centrifugal Filters (30 kDa MWCO). 700 µL of the SL1 buffer of the NucleoSpin Soil DNA extraction kit (Macherey-Nagel) was added to the resulting concentrated supernatant and the cell pellet and DNA extraction was carried out according to the manufacturer’s instructions. Primers (Circ19F 5’-TCT CAC TAA GCA AAA CCC TTA CTT and Circ349R 5’-ACC TCC GTT TAA TAA GAT TTT TCG TGA) were designed to point outwards of both end of the integrated virus, allowing a product detection only on the circularized genome. The qPCR reagent concentrations and cycling conditions were the same as those used for lokiarchaeal 16S rRNA gene detection^12^ with an annealing temperature of 60 °C. Amplified fragments from B36 enrichments DNA were used as standards and the efficiencies of these reactions varied from 90% to 95%, with R^2^ value of > 0.99. Primer specificity was confirmed through Sanger sequencing of the products using the services of Eurofins.

### Virus identification and assembly

Quality trimmed paired-end reads were assembled using metaplasmidSPAdes^59^ and MetaviralSPAdes^60^ to obtain extrachromosomal elements present in our enrichment. VirSorter v2.2.4^61^ was used to identify potential single-contig viral genomes. Putative viral genomes were mapped against the *Ca.* L. ossiferum B36 genome using minimap2 v2.26^62^. The high coverage provirus region within the *Ca.* L. ossiferum B36 genome was identified by mapping back the reads to *Ca.* L. ossiferum B36 genome using bowtie2^63^. Mapped reads were converted to BAM format using SAMtools^64^ and visualized in Geneious Prime® 2022.2.2.

### Induction of viral particle production

In all experimental setups testing the influence of different conditions on viral particle production, 550 mL batch cultures in the exponential growth phase were subsampled (50 mL per replicate) in technical triplicates and exposed to one of the following conditions: introduction of 5%, 10%, or 20% atmospheric air into the gas phase; cold shock (incubation at 4 °C); heat shock (incubation at 30 °C); or supplementation with an N₂:H₂ mixture (75:25%) in the gas phase. Viral particles were quantified in both the cell pellet and the culture supernatant using qPCR (as described above) after 8 h, 48 h, and 7 days in the initial experiments, and after 7, 14, and 21 days in subsequent experiments.

### Isolation and purification of virus particles and sequencing

Cells from combined cultures (about 300 mL for first biological replicate, about 700 mL for second biological replicate) were removed by centrifugation (4 500 × *g* for 45 min) and the supernantant was filtered through 0.45 µm filters to remove remaining cellular contamination. The supernatant was concentrated with 100 kDa filters to 1 mL (Sartorius Vivaspin ® Turbo 15 RC 100 kDa), and transferred on top of CsCl-step gradient [1.7, 1.56, 1.4, 1.2 g/mL] in salt buffer (100 mM Tris pH 7.4, 0.4 M NaCl, 14 mM MgCl_2_, 14 mM MgSO_4_, 9.3 mM KCl, 50mM CaCl)^59^. After ultracentrifugation (288 000 × *g*, 20 h, 4°C) the gradient was separated in 1 mL fractions and viral particles were concentrated and subsequently washed (salt buffer) to remove CsCl residues on Sartorius Vivacon ® 500 (10 000 MWCO) columns to 50-100 µL. 10% of each fraction was stored for microscopy at 4 °C and 90 % of each fraction was used for DNA extraction (Isolate II Genomic DNA Kit, Bioline), and subsequently PCR (Primer forward 5’-AAT AGC AAC AGA AGC AGC C-3’ and reverse 5’-CAT CCA ATA ACC CTC CAA TCC-3’) and sequencing. Library preparation (NEBNext® Ultra™ II) and sequencing (Illumina HiSeq3000, 2 x 150 bp, 1 Gigabase) were performed at the Max Planck-Genome-Centre (Cologne, Germany). Read trimming was performed with Trimmomatic (v.0.36)^48^ (SLIDINGWINDOW:5:20 LEADING:5 TRAILING:5 MINLEN:50 HEADCROP:6). Genome assembly was performed using MetaviralSPAdes^60^ yielding a 16 071 bp circular genome. Reads were mapped to *Ca.* L. ossiferum B36 genome using bowtie2^63^ followed by coverage calculations using SAMtools depth^64^. Raw reads are available in the Sequence Read Archive under BioProject ID: PRJNA1403407.

### Negative staining of viral particles

Out of the concentrated step gradient fraction, as described above, 5 µL were applied onto glow-discharged (20 s, 25 mA; GloQube Plus/Quorum) continuous carbon film-coated copper electron microscopy grid (mesh 300; Electron Microscopy Sciences), incubated for 60 sec, blotted away using filter paper, and subsequently washed by 10 µL of phosphate buffered saline (PBS) (pH 7.4; Gibco), and with 10 µL of 2% Uranyl acetate (AL-Labortechnik & Diagnostik) in dH_2_O (w/v) solution for 60 sec, respectively, blotted away and air dried. Sample was imaged using the TEM Tecnai 10 (Philips) at 80kV using OSIS Megaview G3 (EMSIS) camera at magnification ranging from 39 000x to 135 000x.

### Virus genome analysis

Gene-shared network analyses were performed with vConTACT2^65,66^ against the Viral RefSeq-archaea and Prokaryotic databases v.211, with the option —rel-mode BLASTP and other default parameters. Network results were displayed using Cytoscape^67^.

Structural predictions were performed with AlphalFold 2^28,29^ with default parameters for all generated models. Protein structures were visualized with open source PyMOL Molecular Graphics System, Version 2.2.0^68^.

### Phylogenomic trees and defense system evolution analysis

The defense systems were identified by integrating results from both DefenseFinder v.2.0.0^69^ and PADLOC v.2.0.0^34^, focusing exclusively on defense systems shared between the B35 and B36 strains. Both protein family trees and the Asgard species tree were reconstructed using a standardized maximum-likelihood approach: sequences were aligned using Clustal Omega v.1.2.4^70^ (--max-guidetree-iterations=100 --max-hmm-iterations=100 --iter=100 --output-order=tree-order), low-quality regions were removed with TrimAL v.1.4.rev15^71^ (-gt 0.95 -cons 50), and phylogenetic trees were inferred using IQ-TREE2 v.2.2.2.6 with optimal substitution models selected via ModelFinder^72^. The species tree was constructed from archaeal marker genes identified by GTDB-Tk v.2.4.1^73^. To enable direct comparison with the Asgard species tree, gene trees containing multiple copies per species were collapsed to the species level by replacing multiple tips with their most recent common ancestor (MRCA), ensuring single-tip representation for each species. Topological congruence between gene and species trees was then quantified using Robinson-Foulds (RF) distances, with statistical significance assessed through permutation tests (n = 9,999). In these tests, the mapping between gene copies and species identities was randomized, and for each permutation, a species-level gene tree was reconstructed using neighbor-joining from patristic distances before recalculating RF distance to the species tree. Tree handling and RF calculations were performed in R v.4.5.0^74^ using the ape and phangorn packages. Observed RF distances significantly smaller than the null distribution (p < 0.05) were interpreted as evidence for vertical inheritance and co-evolution, while incongruent topologies suggested horizontal gene transfer or lineage-specific acquisition events.

### Cryo-electron microscopy sample preparation

Loki culture was aseptically removed from the cultivation flasks using a syringe (5 mL; B.Braun) with a needle (0.8×40 mm; B.Braun), 4 mL of culture was transferred into 5 mL tubes with screw cap (Eppendorf), overlaid with N_2_:CO_2_ (80:20) atmosphere (Westfalen). The sample was kept for overnight sedimentation in an incubator (myTemp Mini; Benchmark Scientific) at 20° C. The next day, the supernatant was removed and the sample was gently mixed in the remaining ∼50 μL of media. The sample was again overlaid with N_2_:CO_2_ (80:20) atmosphere and kept at 20° C until vitrification. For vitrification, 4 μL of the sample was applied onto glow-discharged (60 s, 25 mA; GloQube Plus/Quorum) holey carbon film-coated copper electron microscopy grids (R2/2; mesh 200; Quantifoil), and backside-blotted for 4 s using the EM GP2 (Leica) maintained at 90% humidity and 20 °C chamber conditions before being plunge-frozen into liquid ethane. Grids were clipped into Autogrids (NanoSoft) and stored in liquid nitrogen until imaging.

### Cryo-electron tomography data acquisition and processing

Tilt-series were acquired using a 300 kV cryo-TEM Krios G3i (Thermo Fisher Scientific) equipped with a BioQuantum post-column energy filter and a K3 direct detector camera (Gatan) operated at 26 000x nominal magnification resulting in pixel size of 3.37 Å at positions selected after extensive screening using polygon montage option in SerialEM v4.2.4 software (Mastronarde 2005). Acquisition was performed using the PACE-tomo plugin v1.9.2 (Eisenstein 2022) within the SerialEM v4.2.4 software. The cumulative dose of 180 e/A^2 was equally divided into 61 tilts in a dose-symmetric acquisition scheme (Hagen 2017) ranging from −60° to 60° with a 2° increment. The tilt series were acquired at a defocus ranging from −3 μm to −5 μm. Individual tilts were subjected to gain correction, frame alignment, defocus estimation, contrast transfer function (CTF) correction, and split into odd and even in WarpTools v2.0.0 (Tegunov 2019). Assembled and 4x binned tilt-series were subjected to patch tracking alignment in IMOD v5.1 (Kremer 1996). Poor-quality tilts or tilt-series were removed after manual check based on extensive tilts shift or low signal-to-noise (SNR) ratio due to sample thickness. Tilt-series at 4x binning corresponding to pixel size of 13.48 Å were subjected to tomogram reconstruction implemented in WarpTools. Reconstructed tomograms were denoised in Cryo-CARE v0.3.0 (Buchholz 2019) using a model trained on the same dataset. Tomograms were visualized using 3dmod from IMOD.

### Scanning electron microscopy sample preparation and imaging

Overnight-settled Loki samples, as described in the cryo-electron microscopy sample preparation section, were resuspended in last remaining ∼200 μL of media, and applied onto poly-L-Lysine treated (0.01% for 10 min; Merck) circular glass slides (⌀12mm; Epredia) placed inside a 24-well plate (Techno Plastic Products), overlayed with N_2_:CO_2_ (80:20) atmosphere and incubated for 1 hour at 20° Celsius. Subsequently, media was aspirated and cells were fixed by adding 200 μL of 2.5% glutaraldehyde (Electron Microscopy Sciences) in Loki media for 30 minutes. Cells were then washed 2x with 500 μL of dH_2_O for 20 minutes, and dehydrated in an increasing ethanol (Honeywell) series in dH_2_O - 30%, 50%, 70%, 90%; for 20 minutes each, followed by a final 2x wash in 100% ethanol. Cells were then subjected to critical point drying in EM CPD300 (Leica Microsystems), and sputtered with 5 nm gold layer in EM ACE600 (Leica Microsystems). Samples were imaged using the field emission SEM Merlin VP compact (Carl Zeiss) operated at 2 kV equipped with an Everhart-Thornley secondary electron detector (HE-SE2) at magnifications ranging from 5 000x to 25 000x corresponding to pixel size ranging from ∼11 nm to ∼4 nm.

## Data availability

The *Ca.* L. ossiferum strain B36 genome sequence (accession CP163304) was uploaded to GenBank together with the short and long reads of B36 enrichments under BioProject ID PRJNA1091558, BioSample accession SAMN40527350. Proteomic data of *Ca.* Lokiarchaeum ossiferum B36 enrichment culture have been deposited to the ProteomeXchange Consortium via the PRIDE partner repository with the dataset identifier PXD055577. Two representative cryo-electron tomograms have been deposited in the EMDB under accession codes: EMD-56484 and EMD-56485.

## Acknowledgements

We are grateful to Karin Hager, Sarah Harrer, Mahbod Mousavian, Fatma Baraket, Marlene Thalmann and Sophia Lindorfer for excellent technical help with the cultivation of *Ca*. L. ossiferum strains over the past 6 years. This work was funded by FWF 10.55776/EFP25 to FKMS, FB and CS, FWF project Z437 and ERC Advanced Grant #695192: TACKLE to CS and the Max Planck Society (TAS and SE). This research was supported by the Scientific Service Units (SSU) of ISTA through resources provided by the Imaging & Optics Facility (IOF), the Scientific Computing (SciComp) and the Electron Microscopy Facility (EMF), as well as the Lab Support Facility (LSF). We want to thank Victor-Valentin Hodirnau, Vanessa Zheden and Tommaso Constanzo for support in EM sample preparation and data acquisition.

