## Supplementary Figures 1-8 for "Active virus-host system in a Lokiarchaeon culture"

^3^ Institute of Science and Technology Austria (ISTA), [Klosterneuburg](https://maps.google.com/maps?hl=en&gl=at&um=1&ie=UTF-8&fb=1&sa=X&ftid=0x476d0c21eacfcc8b:0xdade8cc8640428b3), Austria

**Supplementary Tables**

**Supplementary Table 1.** Genomic description of bacterial MAGs from Loki-B35 and Loki-B36 enrichments.

**Supplementary Table 2.** Rearrangement events in Loki-B36 genome.

**Supplementary Table 3.** Unique proteins in Loki-B36 genome compared to Loki-B35.

**Supplementary Table 4.** Annotation of proteins of Loki-B36 with their proteomic expression values from Loki-B36 enrichments.

**Supplementary Table 5.** Conditions tested for triggering replication of the Fyl1 virus.

**Supplementary Table 6.** Structure and function prediction of the Fyl1 virus.

**Supplementary Table 7.** Antiviral defense proteins detected encoded by both strains of *Ca.* L. ossiferum and *P. syntrophicum*.

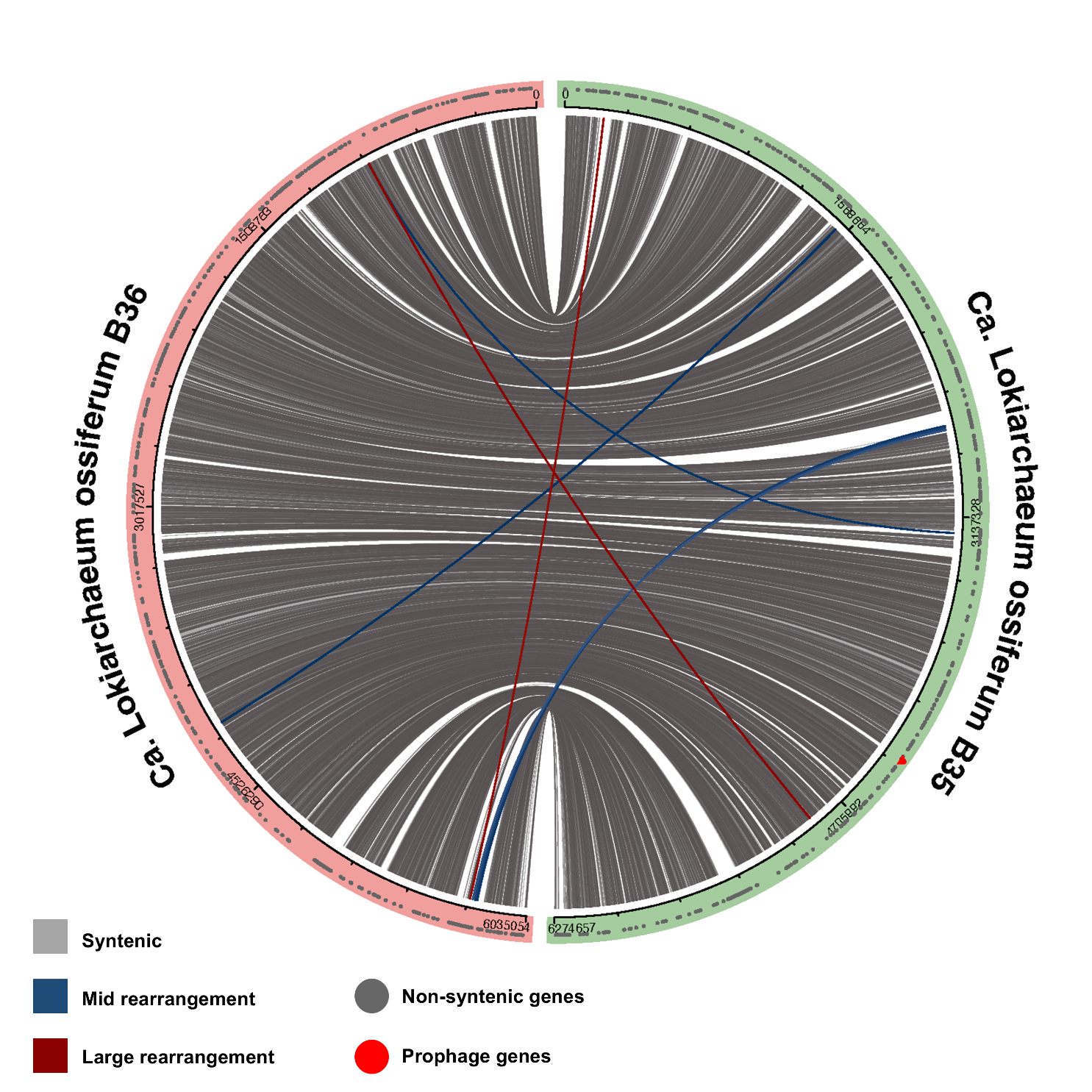

**Supplementary Fig. 1: Genome-level synteny analysis between Loki-B35 and Loki-B36.**
Following the ANI-based comparison strategy, a total of 12 medium-to-long-range genomic rearrangements were identified. A rearrangement is considered large if the distance between the positions in the two genomes is more than 60% of the total genome size. A rearrangement is considered mild if the distance between the positions in the two genomes is between 30-60% of the total genome size.

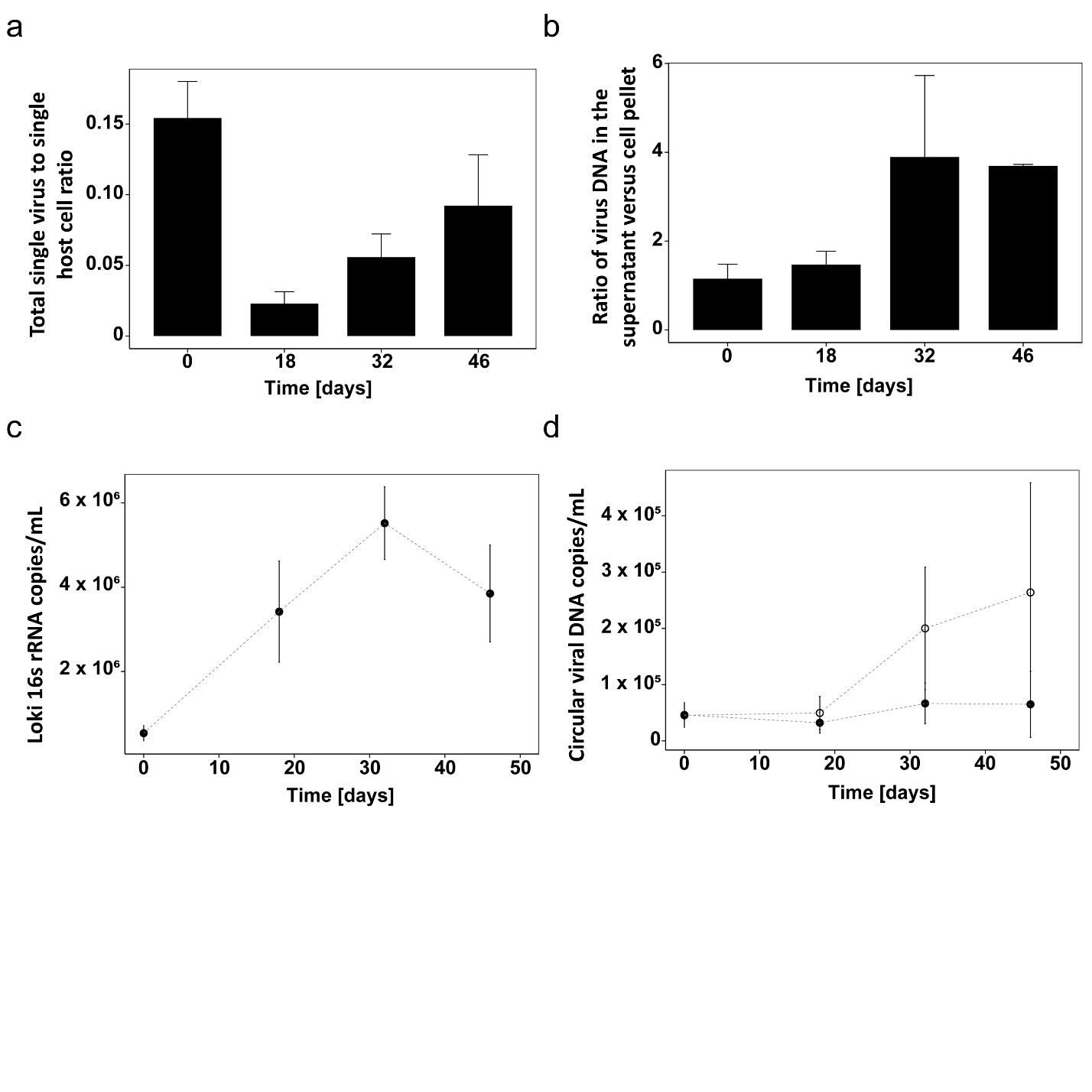

**Supplementary Fig. 2: Host-virus ratios of Loki-B36 and Fyl1. a,** Ratio of total (both in the supernatant and the cell pellet fractions) detected circularized viral genome copies per single Loki-B36 cell calculated via qPCR (see Material and Methods) across lokiarchaeal cell growth. **b,** Ratio of circularized viral genome copies in the supernatant to pellet fractions in the same conditions. **c,** Growth curve of the Loki-B36 cultures used to calculate the ratios above, n=4. **d**, Number of circular virus particles detected in the supernatant (empty circles) and pellet (full circles) fractions in the cultures used to calculate the ratios above, n=4.

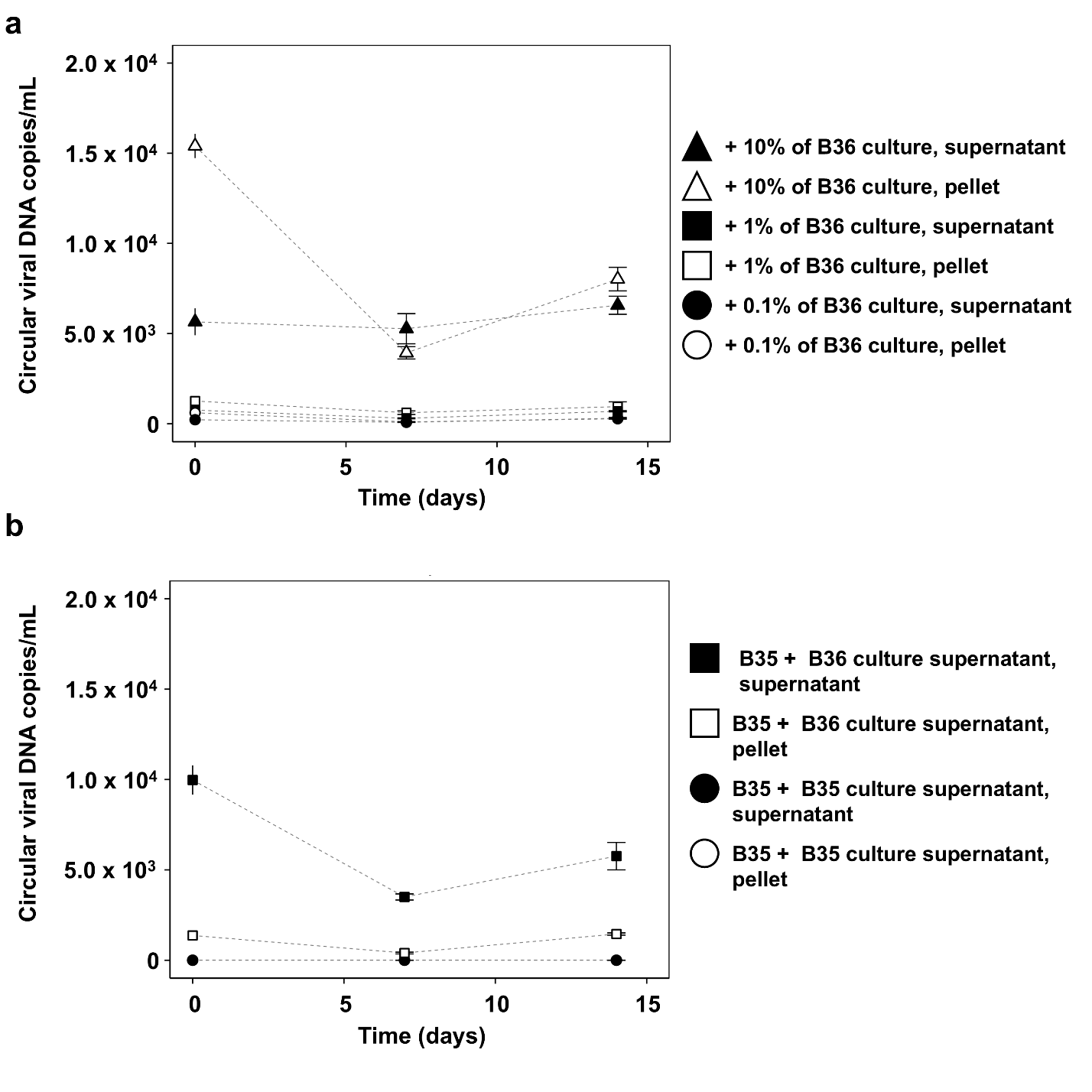

**Supplementary Fig. 3: Infection test of Loki-B35 cells with Fyl1. a,** Number of circular virus particles detected in the supernatant and pellet fractions of Loki-B35 cultures mixed with different amounts of filtered Loki-B36 cultures, n=2. **b,** Number of circular virus particles detecte Loki-B35 (control) culture supernatants, n=2.

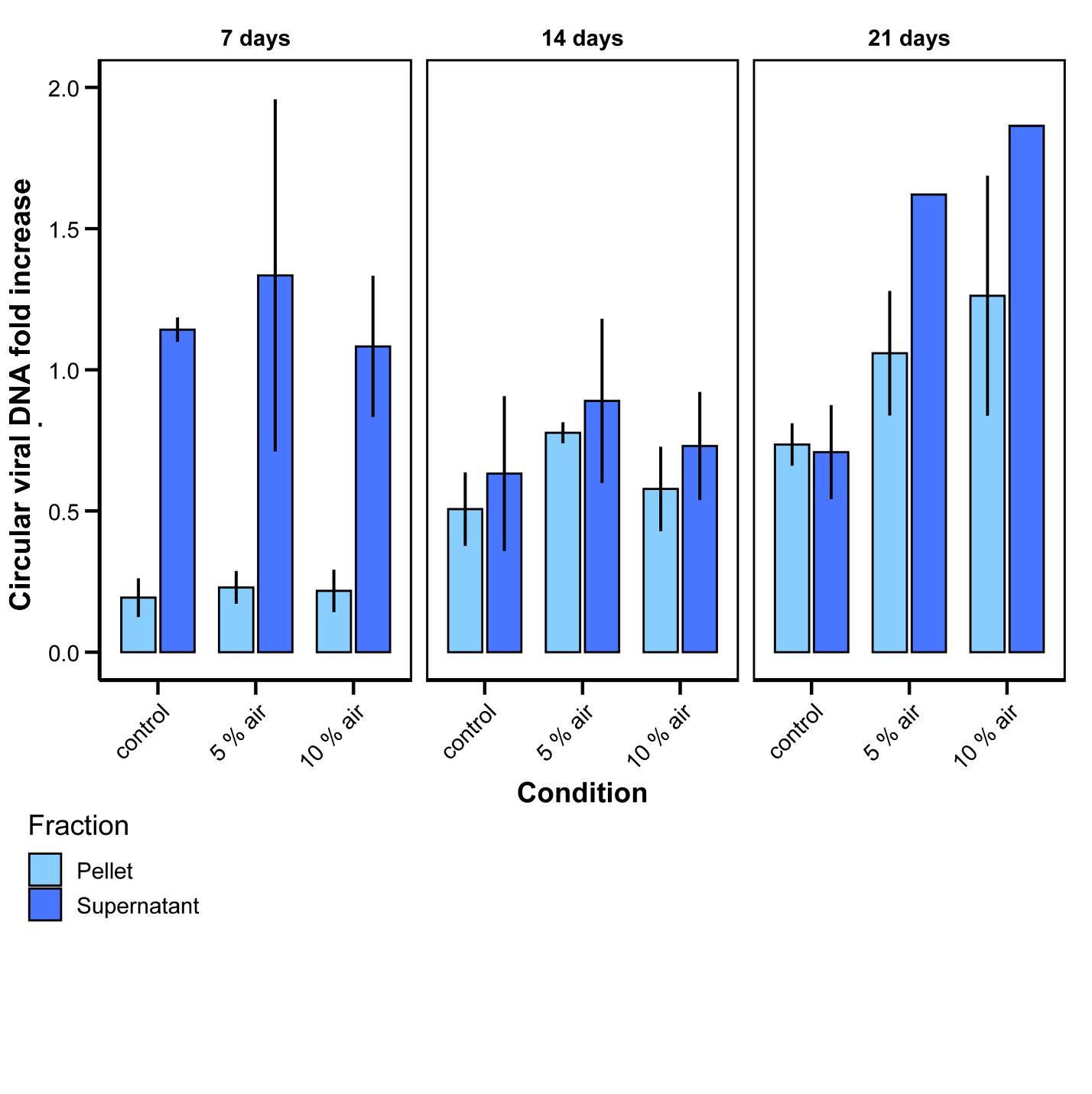

**Supplementary Fig. 4: Induction of Fyl1 with prolonged incubation times.** Influence of different stress conditions on viral particle production in Loki-B36 after prolonged incubation times (n=3). The average increase was calculated by dividing the values obtained from each replicate with an average from the control calculated at time point zero. The number of viral particles was quantified by qPCR in both cell pellets and culture supernatant (see Material and Methods).

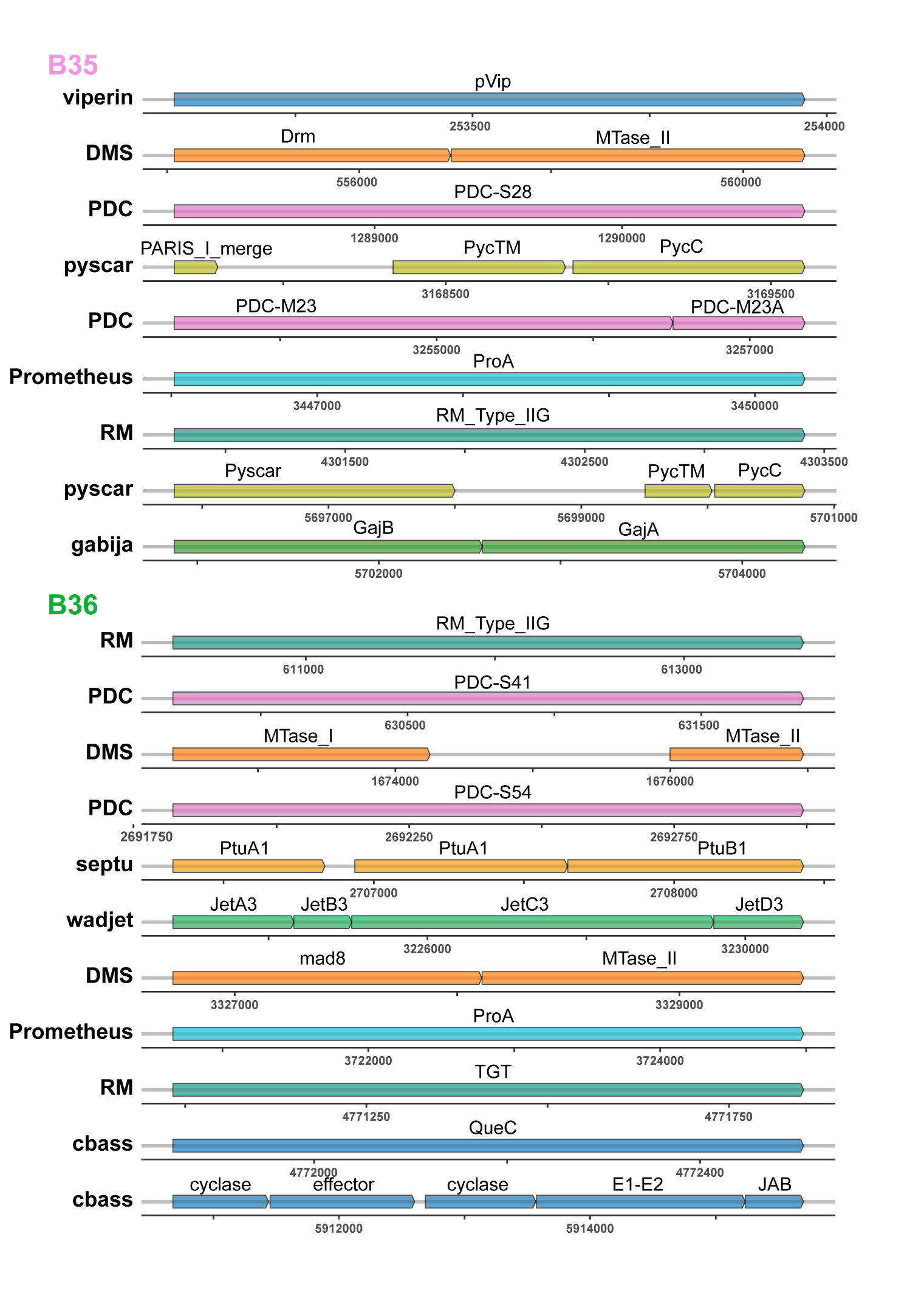
**Supplementary Fig. 5: Comparison of antiviral defense systems of both strains of *Ca.* L. ossiferum.** Numbers indicate the position in the genome.

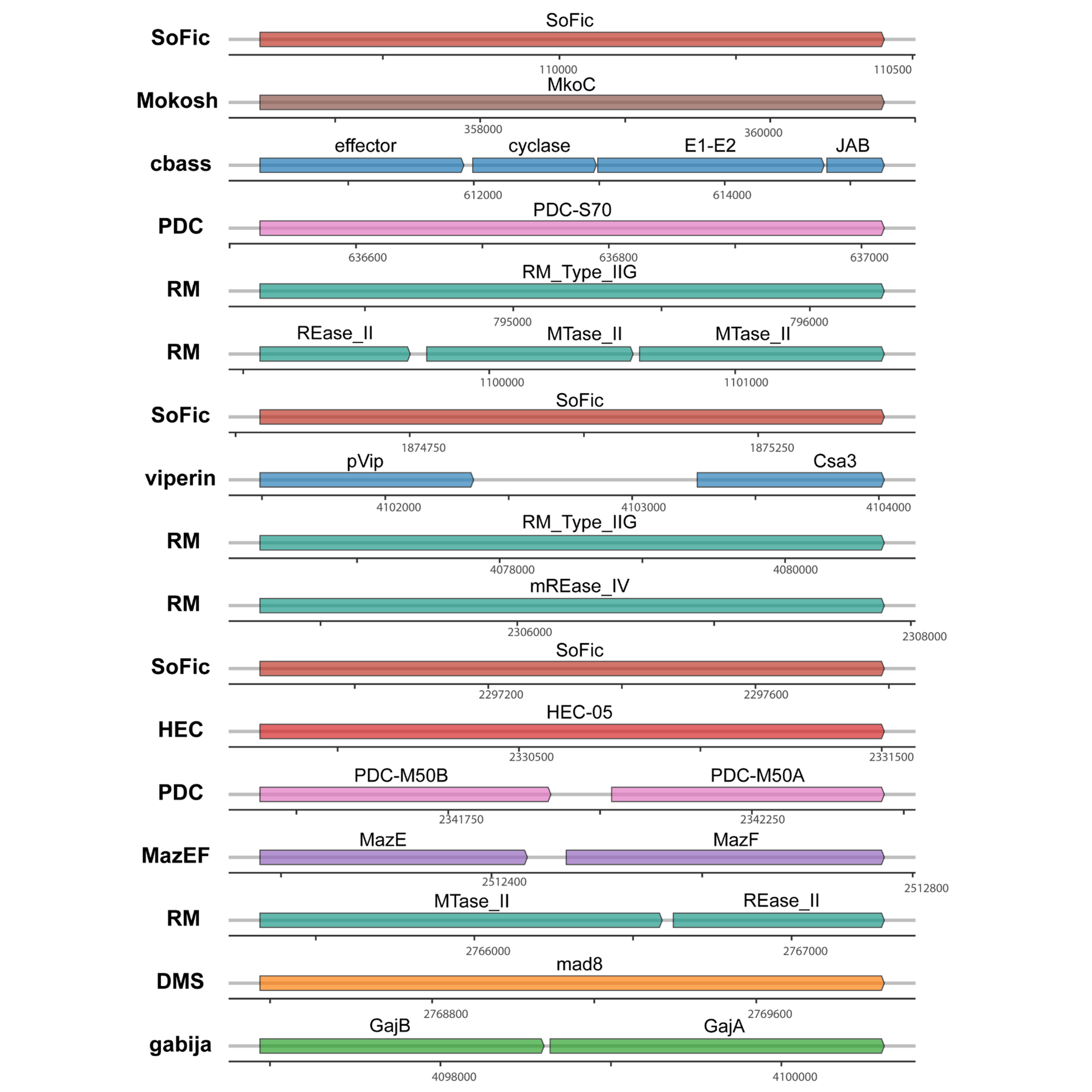
**Supplementary Fig. 6: Antiviral defense systems of *Promethearcheum syntrophicum* MKD1.** Numbers indicate the position in the genome.

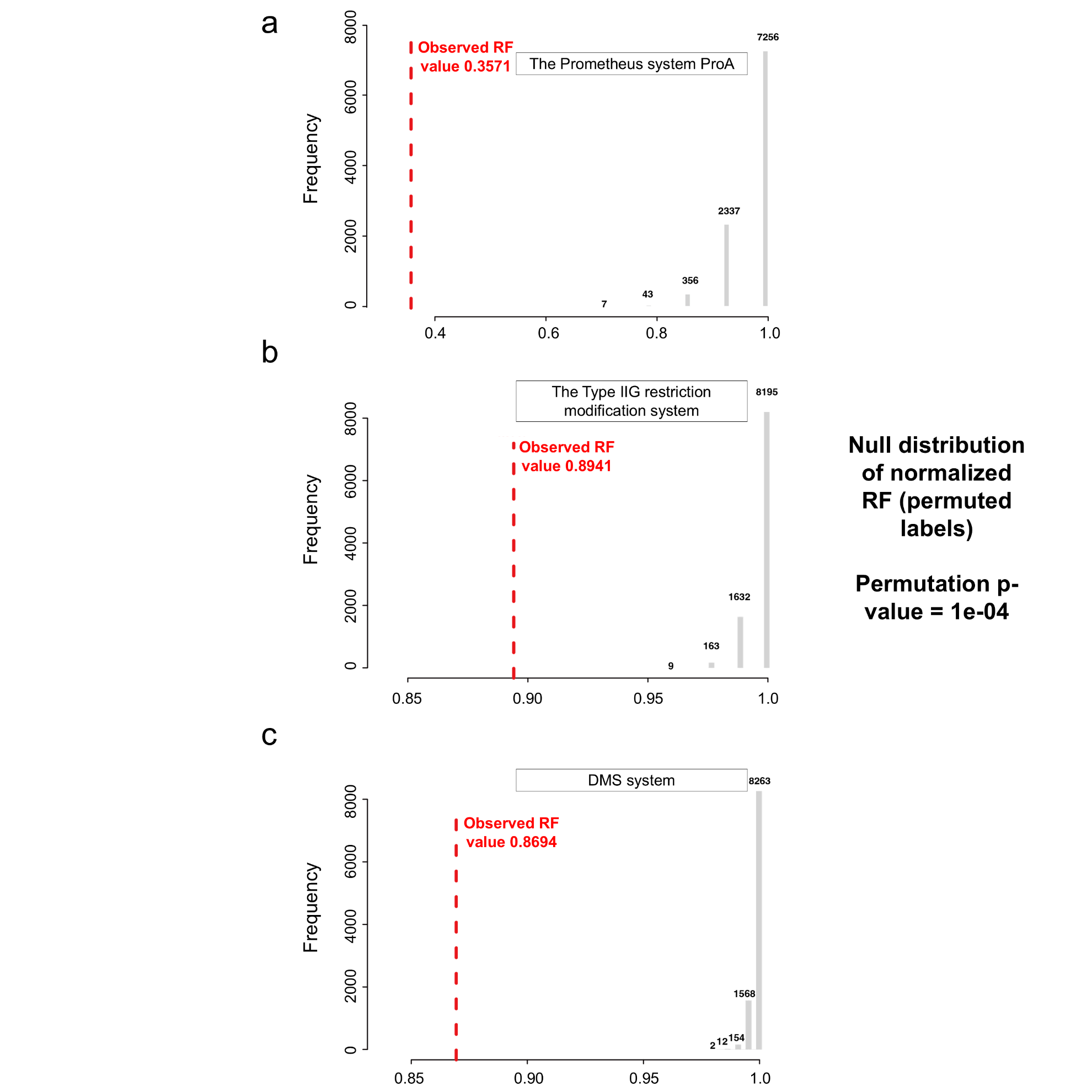

**Supplementary Fig. 7: Topological congruence analysis between the phylogeny of the defense system shared between *Ca.* L. ossiferum B35 and B36 and the Asgard species tree based on the Robinson-Foulds (RF) distances.** Permutation tests for three shared defense system protein families. Each panel shows the null distribution of normalized RF distances (gray histogram) generated from 10,000 permutation tests where the tips of the gene trees were shuffled. The red dashed line indicates the observed RF distance between the actual protein tree and the species tree. Numbers above the histogram bars indicate the frequency of the permuted values in each bin. The permutation p-value is reported at the top of each panel. **a,** The Prometheus system ProA showing significant topological congruence with the species tree (observed RF = 0.3571, p = 1e-04), indicating vertical inheritance and co-evolution with host lineages. **b,** The Type IIG restriction modification system demonstrates strong congruence with the species tree (observed RF = 0.8941, p = 1e-04), supporting vertical transmission. **c,** DMS system exhibiting significant topological similarity to the species tree (observed RF = 0.8694, p = 1e-04), consistent with co-evolution.

These results indicate that all examined shared defense system proteins have likely co-evolved with their host lineages, supporting a scenario of vertical inheritance rather than extensive recent horizontal gene transfer.

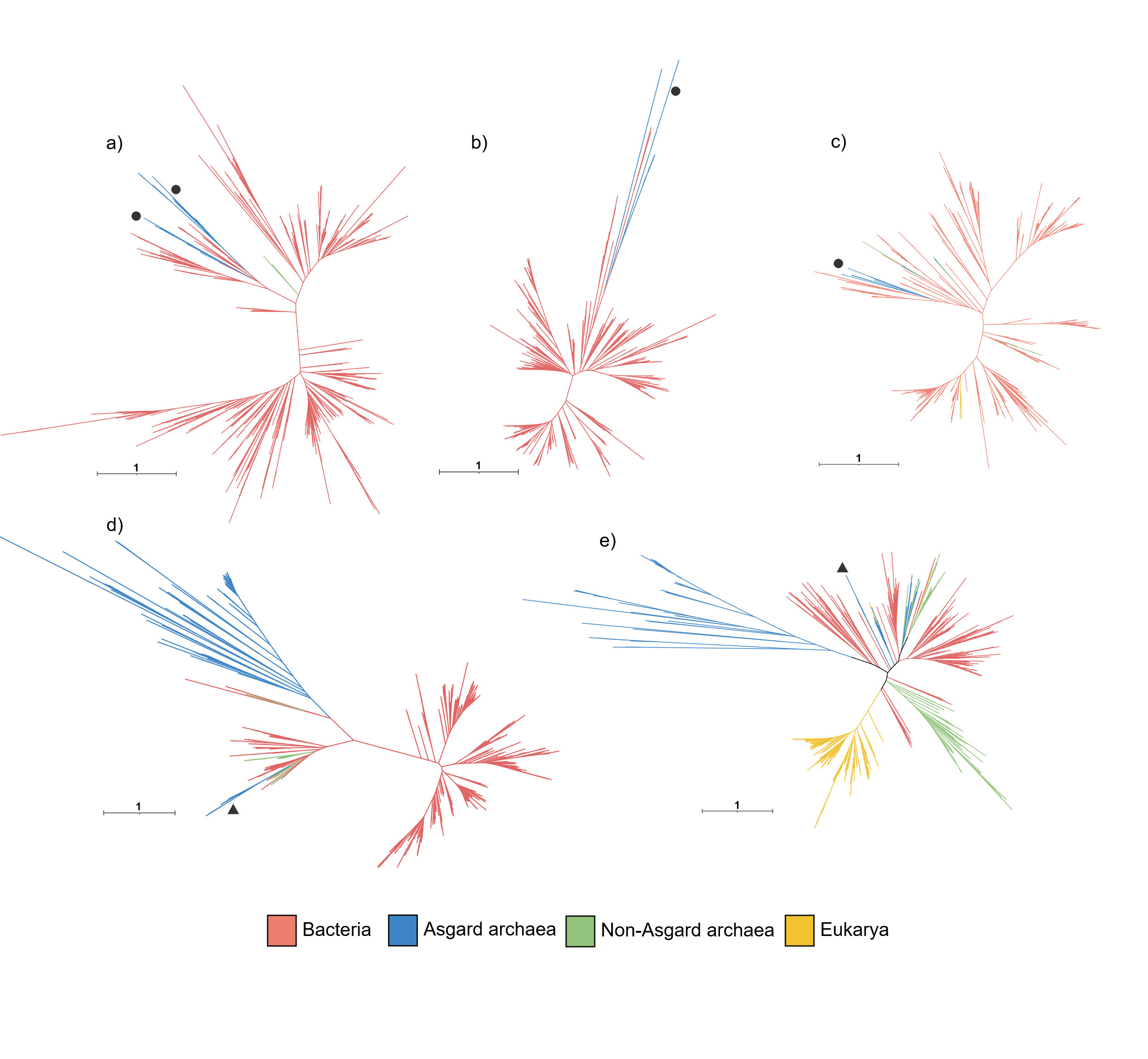

**Supplementary Fig. 8: Maximum-likelihood phylogenetic trees of antiviral defense proteins.** Phylogenies of **a,** cyclases, **b,** effector and **c,** E1-E2 protein of the CBASS system. **d,** Phylogeny of DrmD proteins. **e,** Phylogeny of viperins. The trees were reconstructed using IQtree under the LG+F+G model and 1000 ultrafast bootstrap replicates. Black triangles show the position of *Ca.* L. ossiferum B35 proteins and black circles show the position of *Ca.* L. ossiferum B36 proteins. Bars indicate the tree scales.
